# Solving Paleobotany’s 200-year problem with fluorescence and artificial intelligence

**DOI:** 10.64898/2026.10.12.758007

**Authors:** S. Augusta Maccracken, Richard S. Barclay, Thomas G. Kaye, Matthew J. Butrim, Benjamin Koger, Tyler R. Lyson, Kirk R. Johnson, Ian M. Miller, Samuel S. Robertson, Antoine D. Bercovici, Maria A. Gandolfo, Michael Pittman

## Abstract

Paleobotany has a 200-year problem: leaf fossils are often too poorly preserved to make proper taxonomic identification. Though they are the most abundant, and well-studied macroscopic fossil plant organ, many fossil leaves—including specimens held in museum and university collections—appear to lack the detailed features needed to take crucial next steps, like placement into phylogenetic context, generation of ecological interpretations, or reconstruction of paleoclimate. However, we have discovered that under UV and laser light ∼50% of leaves across a diverse survey of fossil deposits fluoresce in a way that reveals spectacularly detailed leaf venation and gross morphology that is invisible under white light. These ‘Fossil Ciphers’ bring new life into each step of the paleobotanical method, unlocking previously unattainable taxonomic classifications, quantitative morphometrics, and promising new avenues for the incorporation of AI into fossil leaf analyses. With the aid of fluorescence techniques, we can help solve paleobotany’s 200-year problem.

## Introduction

For over 200 years, paleobotanists have been plagued by the inability to place most fossil leaves into proper phylogenetic context. Achieving accurate identification is a common issue for fossils of many clades of plants, but the problem is particularly acute for the leaves of dicotyledonous angiosperms. Their extraordinary diversification in the Early Cretaceous (1, 2), coupled with their variable, plastic, and often convergent morphological characters, makes the classification of their fossil leaves a complex challenge (3). Identifications are based upon leaf architectural characters (i.e. shape, margin, and venation) (4) visible under white light, and under this paradigm the vast majority of fossil angiosperm leaves in research collections lack the morphological fidelity needed for accurate identification. Even for the best-sampled and most intensely studied fossil floras, the number of taxonomically identified species is usually lower than 50% of the known richness of the flora (5), and for more typical floras analyzed using a ‘morphotype’ schema, species-level identifications may be lacking entirely. Lack of identification and misidentification are a problem not only for the field of paleobotany, but also for the molecular phylogenetic research that relies on accurate fossil calibrations (6), as well as any science that leverages our understanding of plant evolution, extinction, diversity, climate, and ecological interactions in the deep past.

Leaf features critical for identification are often obscured by taphonomic processes. Problems arise due to mineral replacement or dissolution of the venation (7), or more commonly through lack of color and/or textural contrast between the leaf margin or venation pattern and the surrounding rock matrix (8, 9). In extreme cases, leaf fossils are morphologically indistinguishable from the surrounding matrix (10); such ‘ghost preservation’ preserves the impression of the fossil but little to no contrast in the compression minerals (Fig. 1A). Similarly, some fossil leaves show little internal contrast among the lamina, veins, and other taxonomically-useful structures (Fig. 1D). These issues of morphological fidelity initially impact specimen collection decisions in the field and go on to affect all subsequent preparation and curation judgement calls in research collections, resulting in collections that do not reflect the recognizable diversity of a fossil deposit. As a result of these taphonomic and sampling issues, many fossil leaf morphotypes and species are erected based on largely incomplete morphological descriptions, or are overlooked altogether. The implications of these preservational shortcomings reverberate throughout the entire paleobotanical method, and have done so since the inception of modern paleobotany more than 200 years ago (11–13). Identifications and analyses of leaf architecture are the bread-and-butter of paleobotany and form the foundation for research into deep-time floral diversity, evolution, and ecology (3), which also provide insights into understanding current and future floral and environmental states. Mitigating, or even circumventing, the influence of poorly preserved leaf fossils on paleobotanical research has the potential to jump-start many crucial discoveries.

**Fig. 1.**
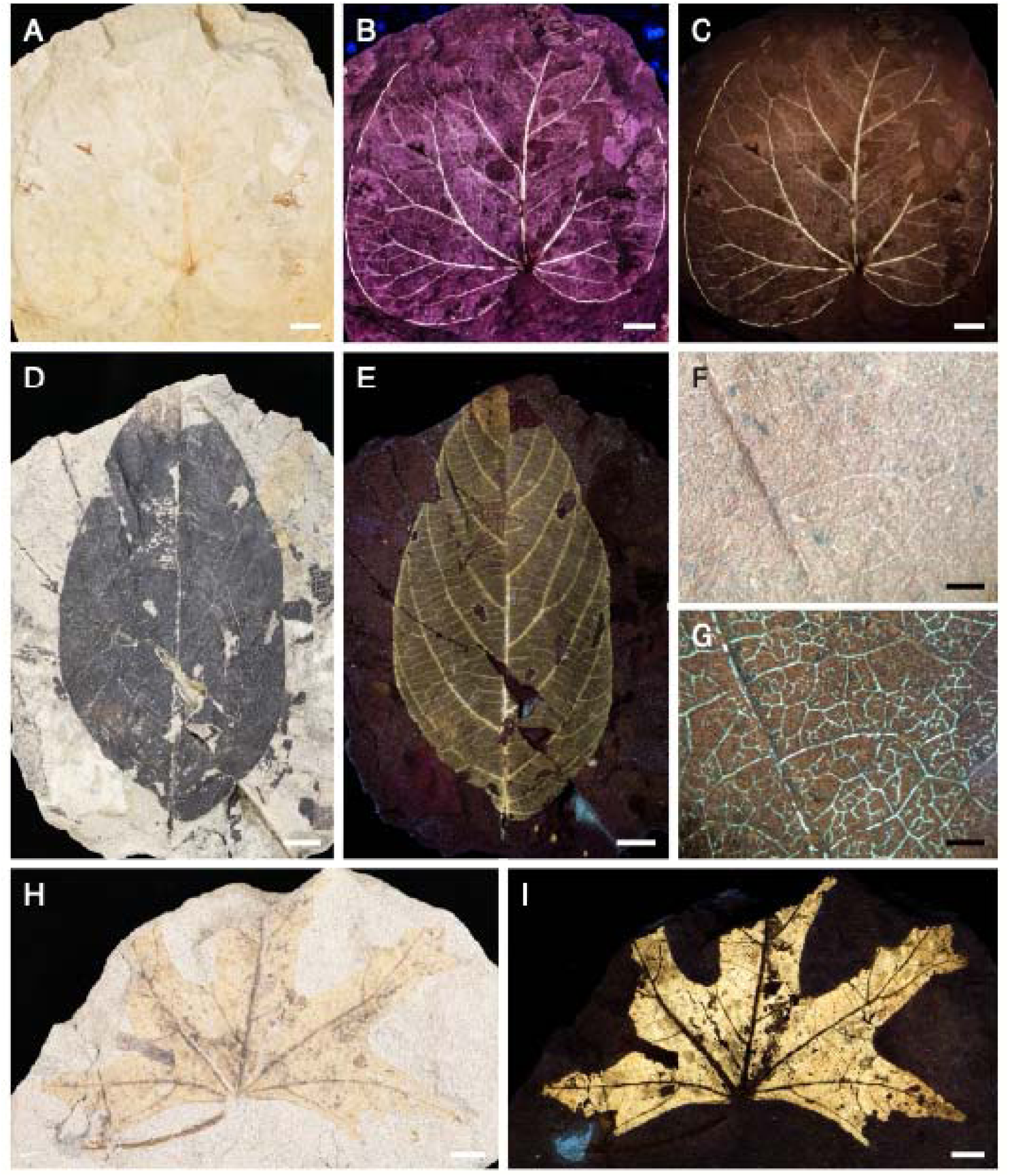
Fossil leaves under incident white light (A,D,F,H), laser fluorescence (B), and UV fluorescence. **(C,E,G,I)**. (**A**–**C**), *Paranymphaea crassifolia* DMNH EPI.20057, Fort Union Fm., Paleocene, DMNH Loc.2200. (**D,E**), Holomorphotype CR72 DMNH EPI.25957 Castle Rock rainforest, Denver Fm., Paleocene, DMNH Loc.2733. (**F,G**). Unidentified dicot DMNH EPI.34578 Denver Fm., Upper Cretaceous, DMNH Loc.2805. (**H,I**). *Cissites panduratus* DMNH EPI.6239, Hell Creek Fm., Upper Cretaceous, DMNH Loc.428. White scale bars = 1.0 cm; black scale bars = 1.0 mm.

Fluorescence occurs when organic and mineral compounds absorb photonic energy which is re-emitted at longer wavelengths, revealing additional detail about composition and morphology normally invisible under white light. Fluorescence imaging has been utilized in paleontological research collections for many decades, such as epifluorescence microscopy of fossil leaf cuticle, and traditionally used shortwave ultraviolet (UV-A) light sources like Wood lamps (mercury vapor or LED based), and more recently, visible violet laser sources. Laser-stimulated fluorescence (LSF) is increasingly utilized in paleontology due to the powerful excitation that the monochromatic and coherent laser light induces on fluorescing compounds, and has led to many exciting new discoveries, particularly allowing vertebrate and invertebrate paleontologists to identify soft-tissue structures (14–20). However, the application of UV fluorescence on macrofloral paleobotanical collections has been limited and that of LSF non-existent.

While it has long been known that fossil leaves can fluoresce, particularly when preserved with cuticle, paleobotanical applications have been restricted to high magnification epifluorescence microscopy (8, 21) and lagerstätten (22–25), where the preservation of leaf characters was never lacking. Our findings, based on either the simple application of a handheld UV flashlight or a portable laser module to ‘poorly’ preserved fossil leaves, show that many fossils in fact preserve key architectural characters, such as their venation pattern and margin details, that were previously invisible under white light. Here we present our discovery of these ‘Fossil Ciphers’, which are fluorescent macrofossils that reveal spectacular leaf architectural preservation under UV or laser light. This finding has the potential to revolutionize our ability to place fossil leaves into phylogenetic context, because it reveals additional venation orders otherwise invisible, maximizes the utility of fossils already in research collections, and aids in developing viable computer vision applications and artificial intelligence to connect leaf venation with biological affinity.

## Results

Application of longwave UV and laser light to Fossil Ciphers reveals leaf architectural characters that were cryptic under white light – particularly for higher-order venation (Fig. 1). We explore the prevalence of Fossil Ciphers within museum collections and confirm their presence across multiple geologic formations in the United States, spanning 300 million years from the Upper Carboniferous Period (Mazon Creek Fm.) through to the Pleistocene Epoch (Swan Lake flora) (Fig. S2). We find that over half the fossil leaves in a Pleistocene locality (DMNH Loc. 95 Swan Lake Willow) and Late Cretaceous locality (DMNH 4150, Kaiparowits Fm.) strongly fluoresce, although the average percentage of strongly fluorescing specimens is ∼18% among all localities (Dataset S1). Dicotyledonous (dicot) foliage is the most frequently encountered plant organ in the survey (n=3,453) and has the highest percentage of fluorescing specimens; 31.68% of all dicot leaves fluoresce strongly and 69.42% at least weakly fluoresce. Many specimens of monocotyledonous (monocot) foliage, gymnosperm foliage, pteridophyte foliage, reproductive organs (seeds, fruits, cones, flowers), and non-reproductive axes (roots, stems, etc.) also fluoresce (Fig. S2). Thus, in general, Fossil Ciphers are far more widespread than expected.

We then quantify the change in visible venation for fossil leaves viewed under white light and fluorescence to determine if and how fluorescence improves our ability to describe fossil specimens. For fossil dicot leaves, the highest leaf vein order (1–8+ vein orders (4)) visible under fluorescence is on average one order higher than that under white light conditions. For example, under white light a fossil leaf might exhibit primary, secondary, and tertiary veins while under fluorescence the primary, secondary, tertiary, and quaternary veins are visible (Fig. 1; Fig. 2). The greatest single increase in visible vein order is for a leaf from the Hell Creek Formation of South Dakota, USA, in which no veins are visible in white light conditions and fluorescence reveals up to the sixth order of venation. While the majority of leaves surveyed show an increase of at least one vein order with fluorescence, some leaves show no improvement and a small number of leaves have a decrease in the highest visible vein order under fluorescence (Fig. 2).

**Fig. 2.**
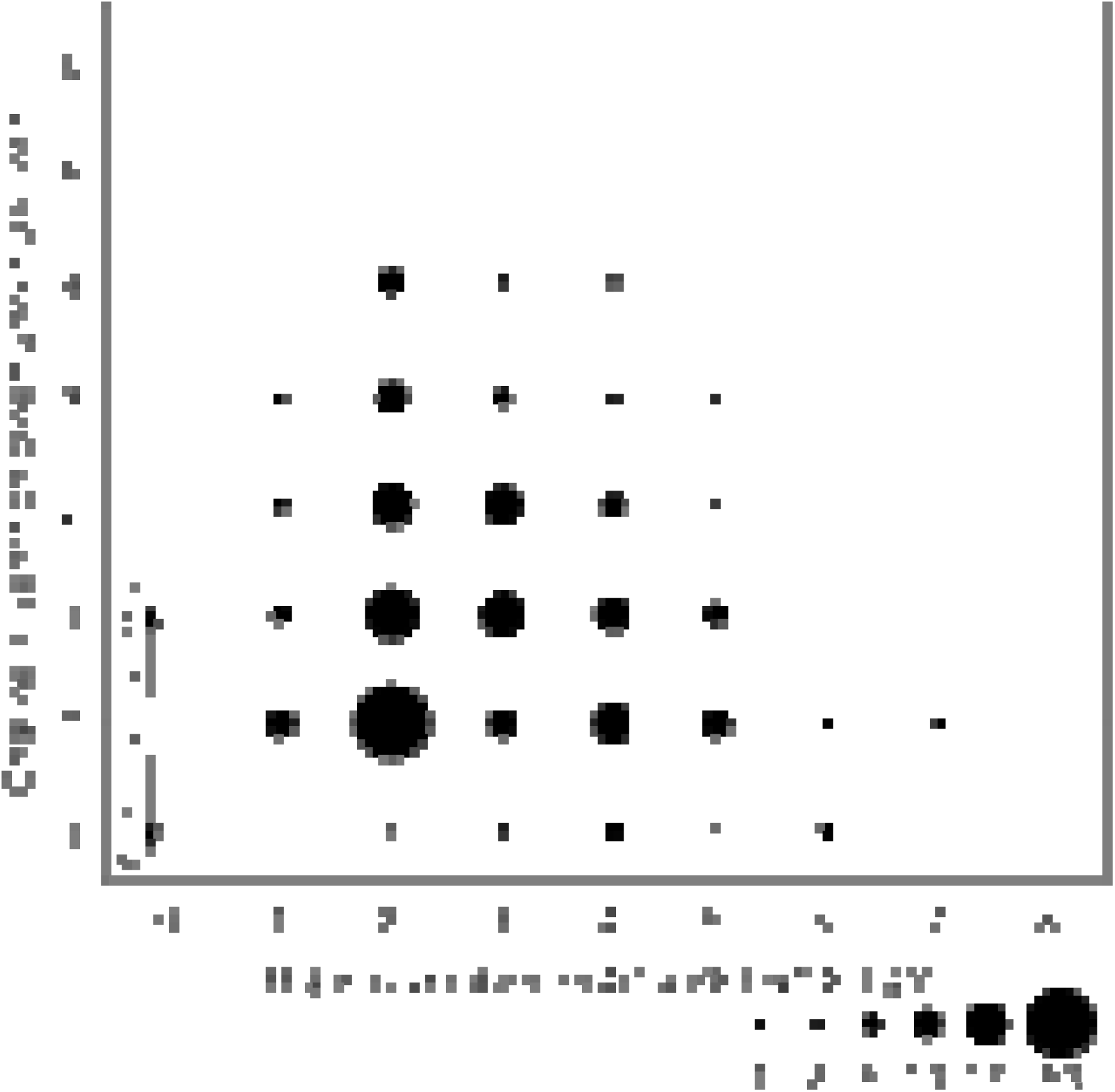
Change in highest detectable vein order for a subsample of fossil dicotyledonous specimens under white light and UV fluorescence. Circle size corresponds to number of specimens (1–50 specimens).

Total vein length is used as a proxy for maximum photosynthetic assimilation rates in fossil taxa, but values can be greatly affected by preservation effects (26). When the quantity of visible venation, measured as total vein length, is analyzed for fossil leaves under white light and fluorescence there is often a substantial increase in the amount of venation visible under fluorescence. The specimens in Figure 1, which represent the most dramatic Fossil Ciphers, have an average of 59% more visible venation under fluorescence compared to that seen under white light conditions. For fossil leaves that have no difference in highest detectable vein order between white light and fluorescence, which serve as a control group, there is an average increase of 28% in visible venation under fluorescence, ranging from 0–67%. The increase in the amount of visible venation for fossil leaves under UV fluorescence also often captures the variability of venation patterns within a leaf compared to that seen under white light conditions, an important component of accurate taxonomic descriptions. Moreover, the particular veins visible under white light and fluorescence may differ, meaning that the fullest complement of identifiable venation is often obtained using both white light and fluorescence to describe a fossil leaf, even in cases where there is a decrease in the visible venation under fluorescence.

Leaf architectural characters of Fossil Ciphers also substantially increase in contrast with various parts of the leaf and surrounding rock matrix, either by fluorescence of veins or lamina, particularly under laser light (Fig. 1B). This quality is successfully leveraged by using a computer vision model to segment diagnostic characters such as leaf margin and venation from multiple specimens (Fig. 3). Fluorescence also assists in the visibility of features not associated with leaf architecture, such as insect or fungal damage (Fig. 4) (27), as well as the preserved body fossils of some scale insects (Sternorrhyncha) (Fig. 4) (also see 28). Additionally, fluorescence may be helpful for detecting chemicals used in preparation and conservation, such as glues, accelerators, and other preservatives (Fig. S1).

**Fig. 3.**
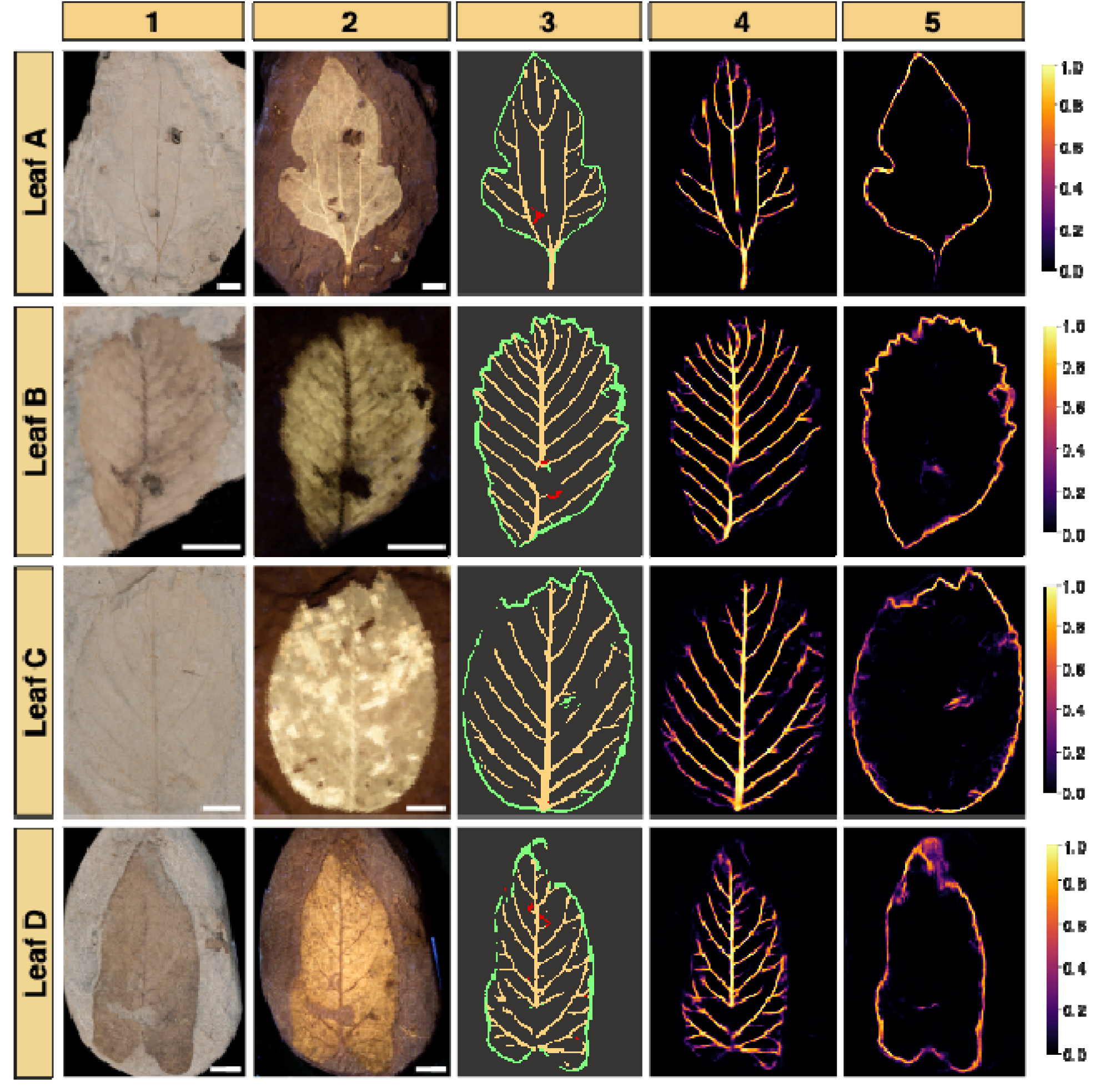
Computer vision model trained on non-fluorescent fossil images and applied to images taken under fluorescence. **(Leaves A,B)**, Laser-Stimulated fluorescence images (Col. 2). (**Leaves C,D**), UV fluorescence images (Col. 2). Automated segmentation of leaf characters (Col. 3): leaf margin (green), primary and secondary venation (yellow), and regions that could cause false positive vein predictions (red). Heatmaps show pixels that the model assigns some probability of being veins (Col. 4) or leaf margin (Col. 5) and their relative confidence (Right). (**Leaf A)**, *Cissites lobata* DMNH EPI.22816, Hell Creek Fm., Upper Cretaceous, DMNH Loc.440. (**Leaf B**), *Paranymphaea crassifolia* DMNH EPI.20057, Fort Union Fm., Paleocene, DMNH Loc.2200. (**Leaf C**), *Dicotylophyllum anomalum* DMNH EPI.77341, Fort Union Fm., Paleocene, DMNH Loc.2612. (**Leaf D**), *Paranymphaea hastata* DMNH EPI.20078, Hell Creek Fm., Upper Cretaceous, DMNH Loc.428. Scale bars = 1.0 cm.

**Fig. 4.**
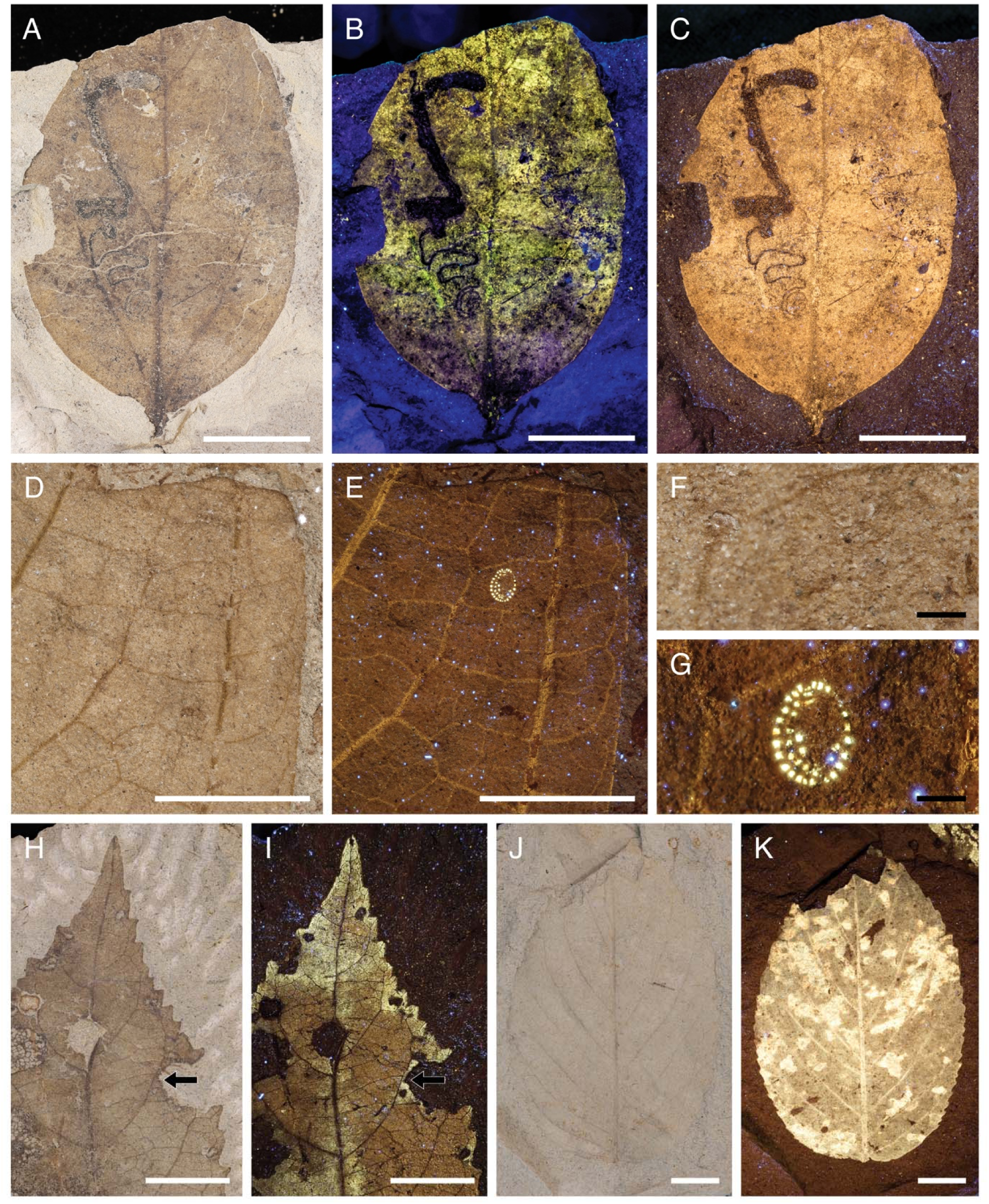
Fossil leaves with insect and pathogen damage under incident white light and fluorescence. (**A,D,F,H,J**), White light fossil leaf images. (**B**), Laser-Stimulated fluorescence image. (**C,E,G,I,K**), UV fluorescence images. (**A,B,C**), Leaf mine on *Catula gettyi* DMNH EPI.47387 Kaiparowits Fm., Upper Cretaceous, DMNH Loc.4150 (36). (**D–G**), Scale insect body fossil on unidentified angiosperm leaf DMNH EPI.45770 from the Dawson Fm., Eocene, DMNH Loc.2484, and closeup of the scale insect (F,G). (**H,I**), Insect-mediated margin feeding damage on an unidentified angiosperm DMNH EPI.47966 from the Kaiparowits Fm., Upper Cretaceous, DMNH Loc.4150. Arrows indicate reactionary scar tissue to the insect damage. (**J,K**), Fossil leaf with pathogen damage on an unidentified angiosperm DMNH EPI.77340 from the Hell Creek Fm., Upper Cretaceous, DMNH Loc.440. White scale bars = 1.0 cm; black scale bars = 1.0 mm.

## Discussion

Fluorescence of plant macrofossils is poised to be an increasingly important tool for describing and therefore identifying fossil leaves. Our extensive survey of fossil collections (4,564 plant organs, including foliage, reproductive structures and non-reproductive axes, the majority of which are dicot leaves) illustrates that Fossil Ciphers are surprisingly common, and that fluorescence increases both the visibility and quantity of the morphological features necessary for accurate identification. These results are especially welcome given the miniscule proportion of formally identified fossil plant taxa compared to that of unidentified specimens in research collections, and especially for highly diverse, dicot-dominated floras of the Late Cretaceous which reflect a decrease in taxonomic confidence with increasing fossil age. For example, the richness of morphotypes and taxonomically identified species from the Hell Creek Formation locality DMNH 566 (5) show the extent of taxonomic identification within an intensely studied fossil flora; of the 24 morphotypes, which include 18 dicots, 15 of the morphotypes are identified to the taxonomic level of species. For these 15 identified taxa, only seven are dicot species (38.9%) that have established systematic affinities (i.e. excluding taxa, such as “*Artocarpus*” *lessigiana*, which need systematic revision). Likewise, the published morphotypic and taxonomic diversity of the Kaiparowits Formation (DMNH Locs. 3642, 3723, 3930) (29) includes 87 total morphotypes, 62 of which are dicot morphotypes. There are eight taxonomic assignments, only three of which represent dicot genera (*Brasenites* sp., *Quereuxia* sp., *Nelumbites* sp.). Notably, seven of the eight taxonomic identifications are aquatic taxa, which are well-known and homogeneous across Campanian age floras, and moreover, no definitive taxonomic identifications are to species level. These floras illustrate the need for increased taxonomic work that will be revolutionized by fluorescence imaging.

Fossil Cipher fluorescence imaged in museum collections, using either laser light or UV (Fig. 1A–C,H–J), can reveal details that will contribute to increasingly complete descriptions of morphotypes and more accurate taxonomic identifications. Our survey suggests that under fluorescence, vast numbers of ‘poorly’ preserved specimens and even floras may yet become identifiable and important in paleobotanical discourse. Our results indicate that portable laser modules provide the strongest and best source for fossil leaf fluorescence, and their ability to change wavelength and intensity of the fluorescence will be instrumental in capturing the range and variation of fluorescence in fossil leaves. Photography using laser light is highly detailed and the color contrast, such as yellow and purple (Fig. 1B), can be especially useful in enhancing and identifying patterns of venation. On the other hand, handheld UV light sources are ideal for broad surveys of curated research collections and are an affordable alternative to portable laser modules, in addition to producing high-quality images. Moreover, bringing an inexpensive battery powered UV LED flashlight into the field to examine specimens *in situ* will help workers make better decisions on which quarries to spend time collecting and which specimens to formally accession in research collections.

These newly discovered well-preserved Fossil Ciphers also hold the potential to push paleobotany squarely into the realm of modern computational analyses. New machine learning techniques that leverage computer vision and artificial intelligence (AI) to extract information from images of fossil leaves have been in development for over a decade (30). Despite the promising nature of these tools, they have not yet been widely applied to actual fossils (31, 32). Perhaps the most significant hurdle to the utilization of these applications on a broad swath of paleobotanical specimens has been the varying quality and composition of fossil leaves (32). Fossil leaf preservation is highly variable in key image attributes, such as color, contrast, and texture, and in object-level attributes, like shape, size, and completeness. Our imaging results show that fluorescence, beyond simply revealing more and finer leaf architectural characters, can create enhanced contrast and a level of preservational consistency that facilitates automated fossil image analysis using AI (Fig. 3), and is often better at doing so than using the corresponding white light images (Fig. S1). Our application of a computer vision model to these fluorescence images resulted in highly successful extraction of leaf macro-features including leaf margin and major venation (Fig. 3), demonstrating that neural networks can consistently identify key leaf features of Fossil Ciphers. Automated segmentation, as shown here, already unlocks a variety of applications–from high throughput morphometric measurements that could be used for paleoclimate reconstructions (e.g. 33) to rapid and unbiased descriptions of leaf architectural traits. Undoubtedly, the most significant findings are yet to come, as our success at segmenting leaf architectural features from Fossil Ciphers heralds the development of new computer models that can bridge our understanding of fossil and modern plant systematics, allowing for more accurate classification (and reclassification) of fossil leaves within the tree of life.

Paleobotanists are aware that fossil leaves with distinct cuticle often fluoresce when using epifluorescence microscopy (24), but the apparent taxon-, locality- and basin-dependence of fluorescence we observe suggests that fluorescence is probably controlled by multiple factors including mineralization that future work will help to elucidate. This fluorescence across fossil floras broadly distributed in time and space raises widespread possibilities for the field of paleobotany and those fields that depend upon fossil leaves, such as paleoentomologists analyzing leaf damage and the remains of fossil insects on leaves (Fig. 4), as well as evolutionary scientists that utilize fossil appearances for calibrations to molecular phylogenies, which rely on accurate taxonomic identifications. In particular, fluorescence of fossil macrofloral remains will become increasingly important for floras deposited during crucial intervals in angiosperm evolution, from the initial rise and radiation of angiosperms in the Early Cretaceous, to the restructuring of floras across major changes in climate (ex. Cretaceous Thermal Maximum (Fig. S2) and Paleocene–Eocene Thermal Maximum), mass extinctions such as that at the Cretaceous–Paleogene boundary (Fig. 1A–C,K,L), and the evolution of new biomes in deep time, such as the evolution of rainforests (Castle Rock; Fig. 1D,E) and floras during the expansion of grasslands.

## Materials and Methods

Fossil leaf fluorescence was imaged using laser light (Laser-Stimulated Fluorescence [LSF]: 405 nm 1W laser diode in a custom-built portable scanning laser module) and long wave ultraviolet light (Yooperlites^©^ C8 Convoy longwave 365 nm filtered UV LED flashlight, with UV safety glasses). A survey of the paleobotanical collections at the Denver Museum of Nature & Science (DMNS) was aimed at analyzing a wide range in temporal and geographic fossil leaf localities. In total, 4,564 elements (i.e. single plant organs such as foliage; reproductive structures including seeds, fruits, flowers, and inflorescences; and non-reproductive axes) from 2,417 specimens were analyzed for fluorescence using handheld UV lights (Table 1), and represent seven geologic formations and 15 fossil localities. The DMNS paleobotany collections are regionally focused, with the majority of fossil floras originating from the Western Interior of North America, as well as temporally focused for deposits from the Late Cretaceous–Eocene, so the survey also reflects these constraints. Localities were also chosen based on sample size, with larger floras (>300 specimens) preferentially selected, although DMNS localities from late Cenozoic deposits were often small (ex. Pleistocene DMNH Loc. 95, n=124 specimens). For the Jurassic Morrison Formation we analyzed seven relatively smaller localities to reach a sufficient sample size. Paleozoic fossil plants at DMNS are generally from historic collections, which are small and have poorly documented locality data, so the survey did not include these specimens. In all, we surveyed collections spanning ∼150 Ma – 12,000 years ago.

**Table 1.** Geologic localities at the Denver Museum of Nature & Science surveyed for fluorescence under UV light.

| Fossil Plant Locality | Geologic Formation | Geographic Location | Epoch & Age | Total Number of Specimens | Total Number of Elements* | Percentage of Strongly Fluorescing Elements | Percentage of Strongly & Weakly Fluorescing Elements |
| --- | --- | --- | --- | --- | --- | --- | --- |
| DMNH Loc. 95<br>Swan Lake Willow | Swan Lake Glacial Sediments | Wyoming, USA | Pleistocene<br>~12,000 years | 21 | 124 | 67.74% | 82.26% |
| DMNH Loc. 90<br>Molloy Ranch | Sucker Creek | Oregon, USA | Miocene<br>~15 Ma | 216 | 231 | 3.46% | 73.16% |
| DMNH Loc. 2484<br>Reagan's Dunn It | Dawson | Colorado, USA | Eocene<br>~55 Ma | 362 | 845 | 17.99% | 57.99% |
| DMNH Loc. 8131<br>Smiley Channel | Denver | Colorado, USA | Paleocene<br>65.81 Ma | 66 | 88 | 15.91% | 45.45% |
| DMNH Loc. 2131<br>Rain Magnet | Denver | Colorado, USA | Paleocene<br>65.98 Ma | 177 | 327 | 11.31% | 59.94% |
| DMNH Loc. 4301<br>Skunk Hunt | Hell Creek | South Dakota, USA | Upper Cretaceous<br>67 Ma | 154 | 419 | 41.76% | 86.97% |
| DMNH Loc. 2805<br>Zebra Beds | Denver | Colorado, USA | Upper Cretaceous<br>~67 Ma | 316 | 514 | 11.87% | 39.49% |
| DMNH Loc. 4150<br>Lost Valley | Kaiparowits | Utah, USA | Upper Cretaceous<br>75.6 Ma | 596 | 1253 | 50.67% | 81.17% |
| DMNH Loc. 3149<br>Bob's Siesta | Morrison | Colorado, USA | Upper Jurassic<br>~150 Ma | 30 | 36 | 22.22% | 55.56% |
| DMNH Loc. 3210<br>Uncle Gorman's Atomic Salad Bar | Morrison | Colorado, USA | Upper Jurassic<br>~150 Ma | 205 | 296 | 6.08% | 16.22% |
| DMNH Loc. 3148<br>Sunspot | Morrison | Colorado, USA | Upper Jurassic<br>~150 Ma | 128 | 172 | 11.63% | 31.40% |
| DMNH Loc. 1506 | Morrison | Colorado, USA | Upper Jurassic<br>~150 Ma | 8 | 9 | 0.00% | 77.78% |
| DMNH Loc. 1507 | Morrison | Colorado, USA | Upper Jurassic<br>~150 Ma | 15 | 35 | 0.00% | 5.71% |
| DMNH Loc. 1505<br>Skunk Quarry | Morrison | Colorado, USA | Upper Jurassic<br>~150 Ma | 80 | 146 | 2.05% | 22.60% |
| DMNH Loc. 2847<br>Ant Hill / Temple Canyon #4 | Morrison | Colorado, USA | Upper Jurassic<br>~150 Ma | 43 | 69 | 5.78% | 53.62% |
| Totals/Averages | — | — | — | 2417 | 4564 | 17.90% | 52.62% |

\*Elements are individual plant organs, such as individual foliage from angiosperms, gymnosperms, ferns, and lycophytes, as well as reproductive organs (seeds, fruits, flowers, inflorescences) and non-reproductive axes (roots, stems, branches).

Fluorescence analyses and imaging were conducted in darkened conditions and following standard safety protocols. Photographs were taken with a Canon EOS 5D DSLR camera fitted with a Canon macro 100 mm macro lens (f/6.3–f/8, 8 s, ISO 400), a StackShot system, and Helicon Focus software (v.8.3); a Nikon DSLR D850 fitted with a Nikon 100 mm macro lens; or an iPhone (12 Pro, iOS v.26). Figures were produced in Adobe InDesign © (v.20.1). LSF images were post-processed across the entire image for equalization, saturation and contrast using Adobe Photoshop (*sensu* 18). No UV fluorescence images were enhanced (i.e. contrast, brightness, exposure) in post-processing. UV light centered on 365 nm produced excellent results (25), but we expect that applying different equipment or filters centered on different wavelengths may be useful for different leaves or characters (18).

The highest order of visible venation was assessed following morphology outlined in the *Manual of Leaf Architecture* (4). Specimens were observed with a stereo microscope (Leica Ivesta 3), first using white light (Motic MLC-150C gooseneck light source), and then in dark conditions using a NIGHTSEA fluorescence viewing system (360–380 nm range, UV-transparent focusing lens, and supplementary filter). 285 leaves from seven localities were included in this analysis. The selected localities spanned the Pleistocene–Late Cretaceous and excluded the Jurassic localities which predate angiosperm remains. Within each of the seven localities, all fluorescent dicots present in one randomly selected drawer were analyzed.

Fossil leaf vein lengths were measured using images of specimens taken under white light and UV excitation. All specimens featured in Figure 1 were selected for this analysis to represent the most dramatic, best-case scenarios for the detection of venation under fluorescence. Additional specimens were then selected to represent cases in which there was no difference in visible vein order between white light and fluorescence (i.e. orders 1:1, 2:2, etc.). This allowed us to test if there was a difference in the amount of venation visible for a particular leaf even if the highest order of visible venation was not different under the two light conditions. We randomly selected two leaves from each of the highest visible venation orders that were the same for white light and fluorescence (orders 1:1–7:7). A total of 14 fossil leaf specimens were included in the analysis.

The visible venation for each image was then traced using an IPad Air (4th gen, v. 15.7) and Apple Pencil for either the entire leaf or a smaller representative section. The images were adjusted for contrast in Photoshop (v.25.11.0) and analyzed in ImageJ2 (v.2.16) using the Ridge Detection plugin (34) to measure the total vein length for each image.

Images of fluorescent fossil leaves were segmented using a DeepLabv3 convolutional neural network model with a Resnet50 backbone (35) trained on a separate dataset of 411 fossil leaf images manually annotated with leaf margin, first order venation, and second order venation masks. This analysis was also conducted on white light images of the same specimens. All data is available on Dryad.

## Acknowledgments

We thank N. Neu-Yagle and K. MacKenzie for their help in developing the UV fluorescing protocol. Thanks to the DMNS Building Ops, especially M. Millard, L. Jones, and S. Murray, volunteer C. Russell, and K. Zubin-Stathopoulos for assistance with data collection, M. Shuley for logistics, and C. Mallinckrodt for assistance with photography.

## Author contributions

Conceptualization & development: SAM, RSB, TGK, MJB, TRL, KRJ, IMM, MP; Data collection & analyses: SAM, RSB, TGK, MJB, BK, KRJ, IMM, SSR, MP; Investigation & writing: SAM, RSB, TGK, MJB, BK, TRL, KRJ, IMM, SSR, ADB, MAG, MP.

## Data and materials availability

All data are available in the main text, the supplementary materials, or Dryad.

## Supplementary Materials

**Fig. S1.**
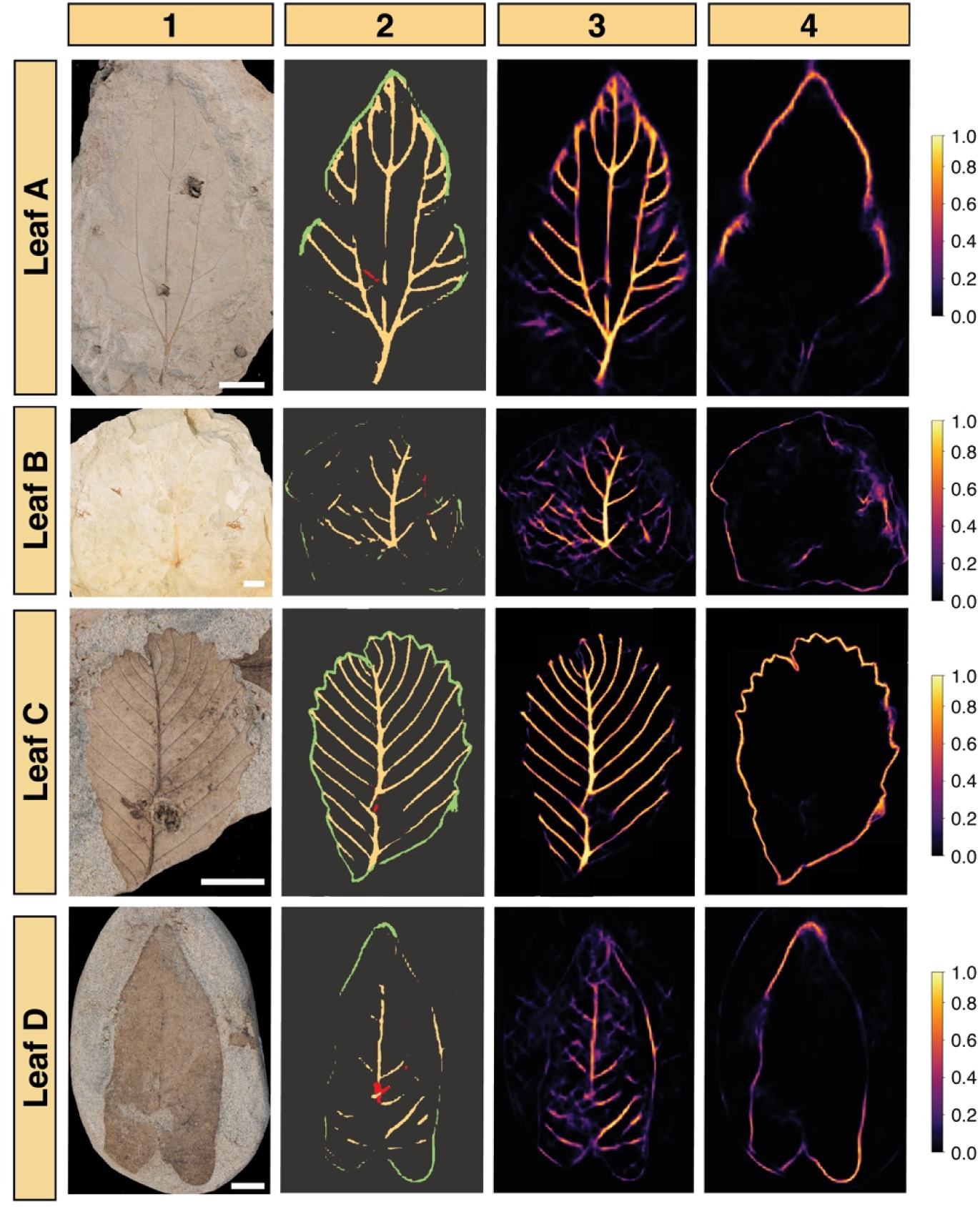
Computer vision model applied to leaves under white light conditions, and corresponding to the analyses in Fig. 3. Automated segmentation of leaf characters (Col. 2): leaf margin (green), primary and secondary venation (yellow), and regions that could cause false positive vein predictions (red). Heatmaps show pixels that the model assigns some probability of being veins (Col. 3) or leaf margin (Col. 4) and their relative confidence (Right). (**Leaf A**), *Cissites lobata* DMNH EPI.22816, Hell Creek Fm., Upper Cretaceous, DMNH Loc.440. (**Leaf B**), *Paranymphaea crassifolia* DMNH EPI.20057, Fort Union Fm., Paleocene, DMNH Loc.2200. (**Leaf C**), *Dicotylophyllum anomalum* DMNH EPI.77341, Fort Union Fm., Paleocene, DMNH Loc.2612. (**Leaf D**), *Paranymphaea hastata* DMNH EPI.20078, Hell Creek Fm., Upper Cretaceous, DMNH Loc.428. Scale bars = 1.0 cm.

**Fig. S2.**
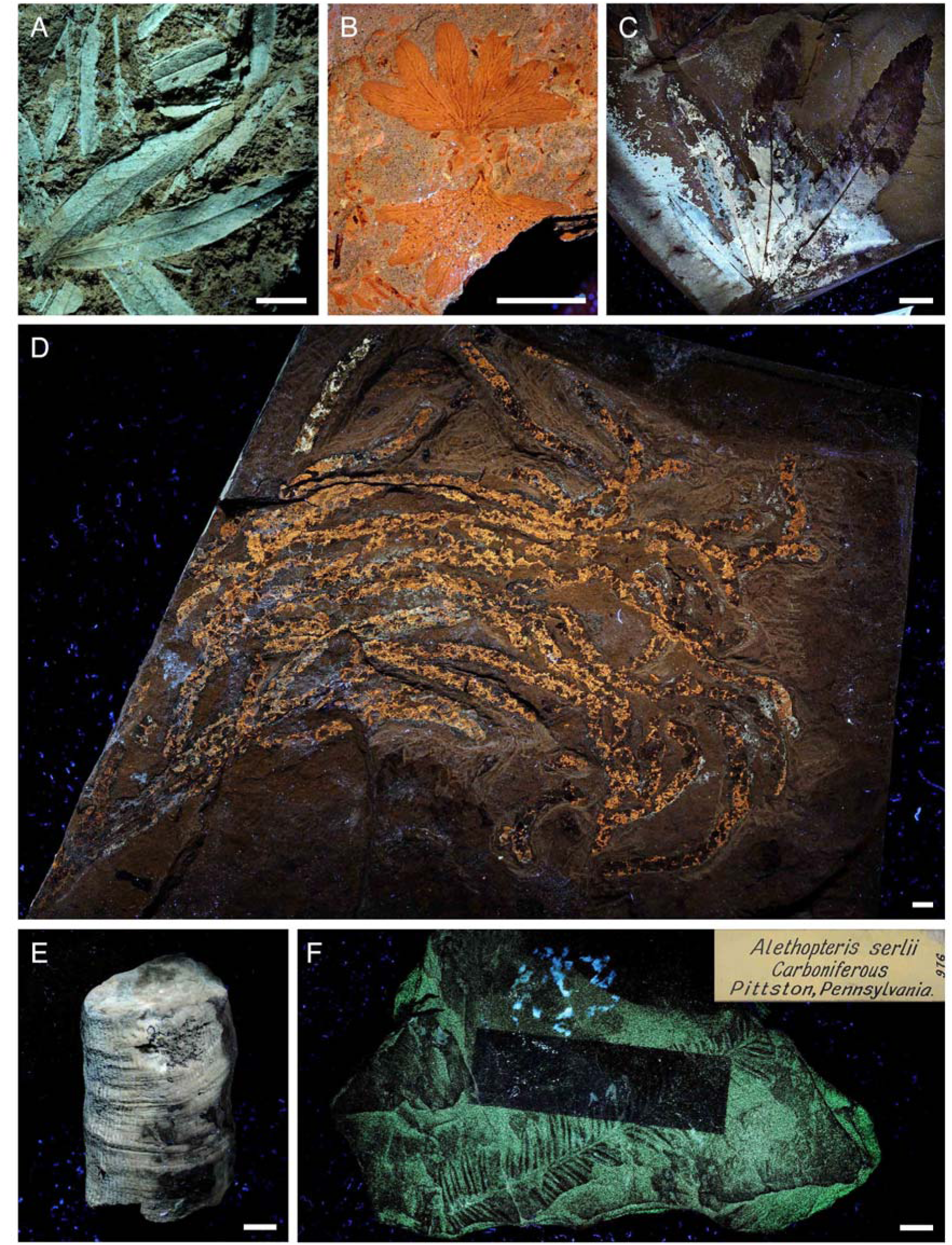
Fossil foliage and stems under UV fluorescence. (**A**), DMNH EPI.60812 Unidentified angiosperm foliage from the Pleistocene Swan Lake glacial sediments of Idaho, USA (DMNH Loc.95). (**B**), DMNH EPI.11226 *Lygodium* sp. fern foliage from the Late Cretaceous Meeteetse Fm. of Wyoming, USA (DMNH Loc.596). (**C**), DMNH EPI.62502 Unidentified angiosperm foliage from the Early Cretaceous Cedar Mountain Fm., Utah, USA (DMNH Loc.). (**D**), DMNH EPI.69577 Gymnosperm material from the Jurassic Morrison Fm. of Colorado, USA (DMNH Loc.3149). (**E**), DMNH EPI.50317 *Calamites* sp. stem pith cast from the Carboniferous Franklin Fm. of Kansas, USA. (**F**), DMNH EPI.976 *Alethopteris serlii* fern fossil from Carboniferous of Pennsylvania, USA. Note the unknown green fluorescing chemical covering the fossil except in the rectangle where the original specimen tag (upper right hand corner of panel F) was placed. Scale bars = 1.0 cm.

Data: S1 to S3 see attached.

